# Loss of chromosome cytoband 13q14.2 orchestrates breast cancer pathogenesis and drug response

**DOI:** 10.1101/2024.06.18.599481

**Authors:** Parastoo Shahrouzi, Youness Azimzade, Wioletta Brankiewicz, Sugandha Bhatia, David Kunke, Derek Richard, Xavier Tekpli, Vessela N. Kristensen, Pascal H.G. Duijf

**Author notes:** **Correspondence:**, &.

## Abstract

Breast cancer (BCa) is a major global health challenge, characterized by chromosomal instability (CIN) and subsequent acquisition of extensive somatic copy number alterations (CNAs). CNAs including amplifications and deletions, significantly influence intra-tumor heterogeneity and the tumor microenvironment (TME). Among these, the loss of chromosome 13q14.2 emerges as a considerable factor in BCa pathogenesis and treatment responses. We provide evidence that this genomic alteration is under positive selective pressure, correlating with poorer patient survival. Furthermore, through multi-omic and *in vitro* analyses, we uncover a dual role of 13q14.2 loss: it confers a survival advantage to tumor cells and modulate the cell cycle and pro-apoptotic pathways in cancer cells, affecting macrophages population in the TME, while paradoxically increasing tumor susceptibility to *BCL2* inhibitors. These findings suggest that targeting 13q14.2 as a biomarker in BCa could enhance the efficacy of existing treatments and offer a new avenue for improving clinical outcomes in BCa.

Figure 1.
Graphical abstract

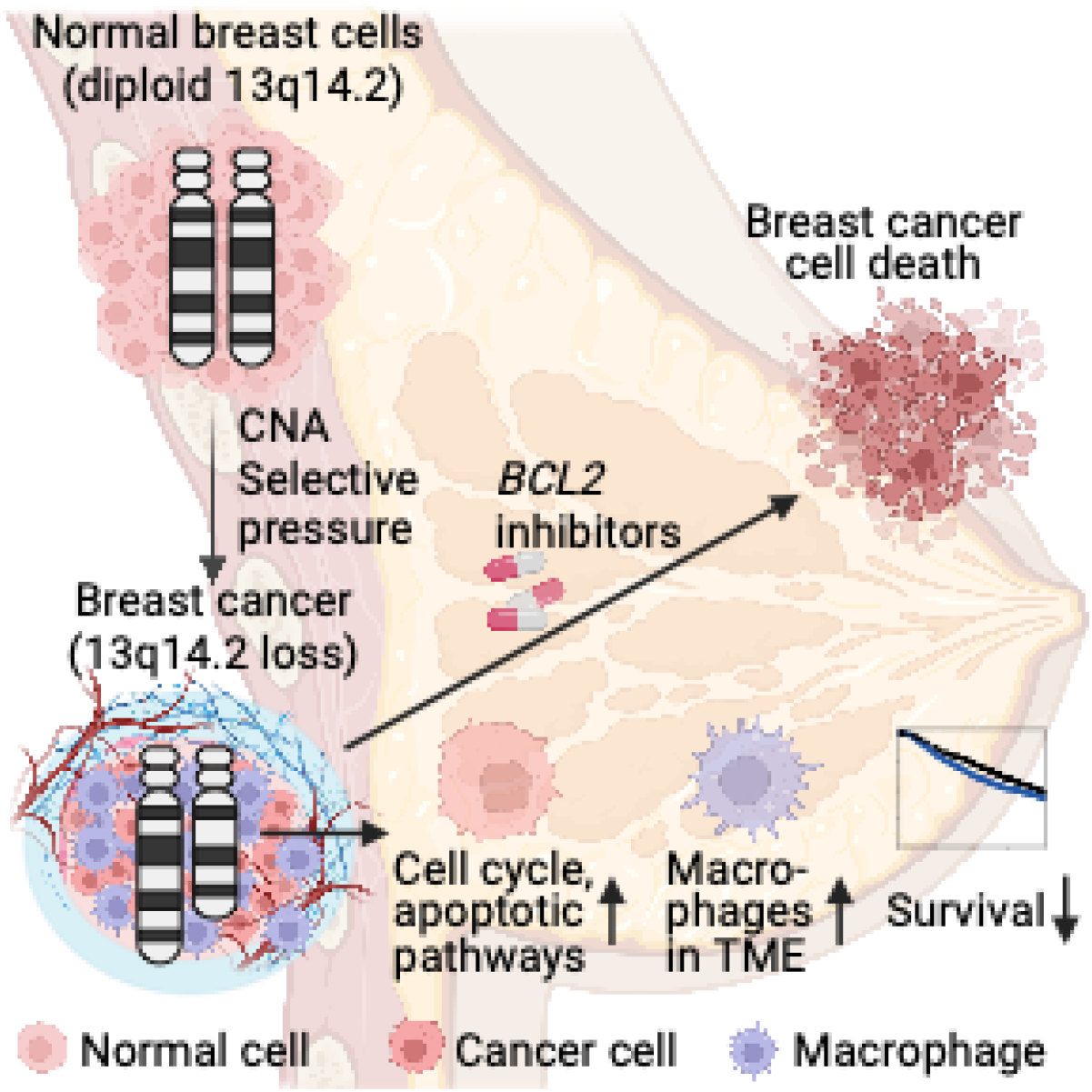

## Introduction

BCa remains a prevalent global concern, affecting approximately one in eight women and one in 726 men throughout their live times (1). While the current classification of BCa – mostly relying on the expression and genomic presence or absence of individual or a group of genes – has advanced therapy selection (2), significant heterogeneity observed within these defined subtypes often results in inadequate treatment response and the development of drug resistance. In addition, chemotherapy is associated with severe side effects in BCa (3). Together, this underscores the critical need for identifying new biomarkers to enable innovative patient stratification and improves treatment approaches.

Tumor heterogeneity in BCa is influenced, in part, by acquisition of genomic abnormalities, such as CNAs (4). Errors in DNA damage repair processes or mitosis cause CIN, leading to acquisition of the CNAs. CNAs involve the gain or amplification and loss or deletion of entire chromosomes and chromo-some arms as well as smaller genomic regions, resulting in aneuploidy and focal CNAs, respectively (5). Many studies have focused on gene-level CNAs, such as those affecting *MYC*, located on chromosome 8q24, widely recognized as the *MYC* amplicon. Our recent research has uncovered a significant functional association between *TRIB1* and *cMYC*, arising through a co-amplification mechanism due to their proximity on the 8q24 locus (6). The co-amplification of other genes in this region alongside *MYC* could have collective effects on tumor biology. This underscores the importance of exploring the broad phenotypic impacts of CNAs beyond individually well-characterized genes.

Chromosomal cytoband 13q14.2 encompasses approximately 50-60 coding and non-coding genes, making it a region of considerable genomic importance. Notably, loss of this region rep-resents the most common CNA observed in chronic lymphocytic leukemia (CLL) (7). In BCa, the focus has been predominantly on mutations or loss of single genes within the 13q14.2 region, such as *Retinoblastoma 1 (RB1)* (8, 9). However, the potential role of 13q14.2 loss as a tumor driver and its impact on the tumor microenvironment (TME) beyond *RB1* loss remains largely unexplored.

While CNAs under positive selection are often drivers of onco-genesis, their presence may not always confer an advantage in the context of anti-cancer therapies. We recently reviewed the dual role of CNAs in drug response, elucidating their potential as drivers of both drug sensitivity and drug resistance (3). The concept of drug susceptibility conferred by specific CNAs is relatively unexplored but holds significant promise for leveraging existing drugs effectively.

The anti-apoptotic *BCL2* protein was first identified in 1984 (10). Thirty-two years later, the Unites States Food and Drug Administration (FDA) approved the *BCL2* inhibitor venetoclax for treating aggressive CLL (11). Overexpression of *BCL2* in 73% of BCas, with 86% in *estrogen receptor-positive (ER+)* tumors, (12) nominates BCa as the first non-hematologic cancer type to be targeted with *BCL2* inhibitors. Currently, the PALVEN clinical trial evaluates the efficacy of venetoclax in com-bination with the cell cycle inhibitor palbociclib and the aromatase inhibitor letrozole in *ER+*, *human epidermal growth fac-tor receptor 2-negative (HER2-)*, and BCL2+ BCa patients (13). While venetoclax has shown promising induction of cell death in hematological malignancies (10), improving treatment response to this drug in BCa may enhance the success of clinical trials in two ways: (1) by utilizing combination therapy to overcome resistance to single therapies and (2) by selecting patients whose cancers are most likely to respond to the drug.

In this study, we hypothesized that loss of 13q14.2 is a significant pathological event in BCa that also influences drug response. To test this, we first investigated the prevalence of 13q14.2 loss in BCas and its association with patient survival. Furthermore, we studied the impact of 13q14.2 loss on cancer cells and their surrounding microenvironment. Finally, we examined the extend, to which 13q14.2 loss affects the sensitivity of BCa cells to venetoclax and other BCL2 inhibitors.

## Methods

### Bioinformatics analyses

GISTIC-threshold-by-gene-based copy number data of the TCGA pan-cancer cohort (ICGC/TCGA, Nature 2020) and the METABRIC breast cancer cohort (Nature 2012 and Nature Communication 2016) were downloaded from cBioPortal (14). The analyses of CNAs, gene expression, protein expression and clinical data in TCGA and METABRIC were performed using *R* programming language (version 4.3.3) and the figures were produced using the *ggplot2 R* package, version 3.5.1.

### Survival analysis

Overall survival analyses were performed by comparing BCa patients with 13q14.2 copy number loss to those without 13q14.2 loss. *Survival R* package (15) was employed. In addition to all patients together, patient survival was also assessed per molecular subtype. Overall survival data were extracted from the TCGA clinical data files and matched to copy number data by patient ID.

### Mutual exclusivity and co-occurrence analyses

For co-occurrence analysis of 13q14.2 loss and top mutated genes in BCa, we first prepared the data by adding a binary column for 13q.14 loss, indicating the presence or absence of the event. Mutations for other genes were also included with genes as columns, samples as rows, and mutations represented as binary (1 for presence, 0 for absence). We then conducted Fisher’s exact test for each gene by looping through each gene, constructing a 2×2 contingency table for that gene with 13q.14 loss. Fisher’s Exact Test returns two values: p-value and odds ratio for each test. The odds ratio indicates the strength and direction of the association; values greater than 1 suggest co-occurrence, while values less than 1 suggest mutual exclusivity. For each alteration, the Direction was calculated using the log2 of odds ratio, multiplied by-log10(p value). Genes with significant p values (<0.05) were kept. Genes with positive-log10(p value) * Direction co-occur with 13q.14 loss, while genes with negative-log10(p value)*Direction are mutually exclusive. The same approach was used to explore co-occurrence and mutual exclusivity of copy number loss of 13q14.2 and copy number loss or gain of other cytobands.

### Differential expression analyses

Differential gene expression analyses were performed using the TGCA breast cancer dataset. Gene expression levels were compared between samples with 13q14.2 loss (n=464) and samples without 13q14.2 loss (n=486). For each gene, gene expression levels were scaled to obtain z-scores per sample. The differences between the median gene expression levels were determined for each gene and adjusted *p* values were calculated using Wilcoxon signed-rank tests followed by Bonferroni-adjustment. Protein expression levels were similarly compared, except that *p* values were adjusted using the Benjamini–Hochberg method (16).

### Pathway enrichment analyses

To identify activated pathways as a consequence of 13q14.2 loss, the top 50 differentially upregulated genes were used and subjected to analyses with MSigDB (17), Reactome (18) and Panther (19). Only upregulated genes were used to avoid potential false discovery of path-ways due to the inclusion of genes located in the 13q14.2 locus that were observed to be preferentially downregulated (Fig. 5a).

***Deconvolution.*** Deconvolution of bulk expression or RNA seq data refers to the computational process of estimating the pro-portions of different cell types that generate the final aggregate measurement. Similar to multiple linear regression model ap-plied to gene expression (microarray or bulk RNA seq) data, it can be formulated as (20):

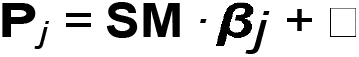

where **P***_j_* is the measured expression/RNA-seq values from tumor sample *j*, **SM** is a Signature Matrix (SM) — that is, a cell-specific expression profile — and β*_j_* is a vector of mix-ing fractions for sample *j*, and represents the noise. Thus, Σ_*k*_ for *P_ij_* being the expression of gene *i* in sample *j*, we have: *P_ij_* = *SM_ik_*β*_kj_* in which *SM_ik_* is the averaged expression of gene *i* in cell type *k* and β*_kj_* is fraction of cell type *k* in sam-ple *j*. Deconvolution has gained increasing attention and a wide range of methods have been developed (21). Here, we utilized a common method known as CIBERSORTx (22). Bulk RNA-seq data of TCGA and GEPs data of METABRIC were down-loaded form cBioPortal. TCGA data were log2-normalized and scaled and METABRIC data were scaled. We estimated cell fractions using either "Imput Cell Fractions" module of CIBER-SORTx webtool with default settings or the Docker version of CIBERSORTx with a similar setting. Following the pipeline we developed for more reliable and reproducible deconvolution (23), we estimate cell fractions using previously created 10 SMs (created using high resolution single-cell RNA sequencing (scRNA-seq) from (24)), and then averaged over all 10 estimates. We also perform an analysis on association of loss of 13q14.2 and fraction of cell types enumerated from spatial omics data. We used imaging mass cytometry data for a subset of METABRIC cohort (25) which were downloaded from the https://zenodo.org/record/5850952 and compare fraction of different cell types.

### Drug response analyses

Drug response analyses of 52 breast cancer cell lines were conducted using the Genomic Drug Sensitivity in Cancer (GDSC) datasets GDSC1 and GDSC2 (26), version 8.4 (June 2022). Genomic features included in the analyses were recurrent mutations and focal copy number alterations, as described (27, 28). A machine learning approach (elastic net regularisation (29)) was applied to identify pharmacogenomic interactions, as determined by effect sizes (Glass’Δ of IC_50_ values) and associated *p* values. These analyses were performed both genome-wide and pertaining to pharmacogenomic interactions involving 13q14.2 loss only. For direct comparisons be-tween cell lines with copy number neutral and loss of 13q14.2, *p* values were determined using non-parametric Mann-Whitney *U* tests. Effect sizes and 95% confidence intervals were determined using point biserial correlation coefficient *r*, with *r* value boundaries between negligible, small, medium and large at 0.10, 0.24 and 0.37, respectively (30).

### Patient-derived xenografts

Drug responses to navitoclax in breast cancer patient-derived xenografts (PDXs) were evaluated using the area under the curve (AUC), as determined in published work (31). Copy number profiling determined by shallow whole-genome sequencing was also described in this study, enabling separation of PDXs into groups with and without 13q14.2 loss.

### Cell culture

Human BCa cell lines MFM223, CAL120 and EVSA-T were purchased from Leibniz-Institut DSMZ-Deutsche Sammlung von Mikroorganismen und Zellkulturen GmbH, who provided the authentication certificate. Human BCa cell lines DU4475, BT474, MCF7, BT20, HCC1187, HCC2218 and HCC1500 were purchased from American Type Culture Col-lection (ATCC). None of the cell lines used in this study were found in the database of commonly misidentified cell lines maintained by the International Cell Line Authentication Committee and NCBI Bio sample. All cell lines were routinely monitored for mycoplasma contamination (MycoAlertTM mycoplasma detection kit, Lonza Cat#LT07-318). MFM223 and EVSA-T cell lines were maintained in EMEM (ThermoFisher Cat#31095029) media. BT20 was maintained in DMEM (ThermoFisher cat#11965084). DU4475, BT474, MCF7, HCC1187 cell lines were maintained in RPMI-1640 media (ThermoFisher Cat# A1049101). Regardless of the media, all cell lines were supplemented with 10% FBS (ThermoFisher Cat#10500064) and 1% penicillin–streptomycin (Gibco; 10,000 U/mL). The PMC42-ET cell line was generated with BCa patient consent and obtained from a pleural effusion by Dr. Robert Whitehead (Ludwig Institute for Cancer Research, Melbourne, Australia), under appropriate institutional ethics clearance (Institutional Review Board of the Peter MacCallum Hospital, Melbourne)(32, 33) The PMC42-LA subline was a spontaneous derivative sub-line from the parental PMC42-ET cells by Dr. Leigh Ackland (Deakin University, Melbourne, Australia) (34). and had been well characterized as a spontaneous and unique MET (mesenchymal-epithelial transition) model system!,which was observed to have more epithelial features than the parental mesenchymal PMC42-ET (35, 36).The whole multi-omics profiling was also well characterized recently including G-banding karyotype, whole exome sequencing and incorporating transcriptomes and proteomics data from this cell model system (37). PMC42 cell lines were maintained in Dulbecco’s modified Eagle’s medium (DMEM) containing glucose (4.5 g/L), L-Glutamine (0.5 g/L) and sodium pyruvate (0.1 g/L) (GibcoTM, Thermo, Catalog number—11885084), and supplemented with 10% fetal bovine serum (FBS; GibcoTM, Thermo, Victoria, Australia) and antibiotics, penicillin and streptomycin (GibcoTM, Life Technologies Catalog number—15140122).

### Drug treatment and cell viability assay

Adherent BCa cell lines were seeded into 96-well plates based on optimal growth rates determined empirically for each line (15% confluency in day 1). 24 hours after plating, cells were dosed with 9 series (3-fold series) of drug concentrations (ABT199, Abcam #ab217298) for 72 h. In case of cell lines in suspension, cells were seeded and dosed with the drugs in the same day. CellTiter-Glo® Luminescent Cell Viability Assay (Promega, #G7571) was added (50% v/v) to the cells, and the plates incubated for 10 min prior to luminescent detection in a Victor 2 microplate reader (Perkin Elmer, Waltham, MA, USA) and the signal was quantified at 595 nm. In case of PMC42 cell lines, cells were treated with drugs (ABT199, ABT737 and ABT263 purchased from Sapphire Bioscience, Australia) and the viability was assessed by the resazurin-based Alamar Blue assay (#R7017, Sigma-Aldrich, St. Louis, MO, USA) and the florescence intensity in each well was measured after 90 mins using a bottom-reading florescent plate reader (CLARIOstar Plus, BMG LABTECH) with excitation at 544 nm and emission at 590 nm.

## Genotyping

### Cell line identification

The identities of all BCa cell lines employed in this study were confirmed by short tandem repeat (STR) profiling.

### SNP6 microarray analysis

Genomic DNA from BCa cell lines was extracted using the Qiagen Blood & Cell Culture DNA Mini Kit (cat #13323) and subjected to SNP6 microarray analysis by the Australian Translational Genomics Centre (ATGC) at the Translational Research Institute (TRI), Princess Alexandra Hospital, using Illumina InfiniumTM Global Screening Array (GSA). The microarray data were analyzed using open-access GenomeStudio Software in conjunction with the CNVPartition plug-in (version 3.2.0).

### Shallow whole genome sequencing

Shallow whole genome analysis was performed on the genomic DNA from a BCa cell line at the Genomic Core Facility at the Amsterdam UMC, Netherlands. The data were analyzed and visualized using Ab-solute Copy Number Estimation in Bioconductor.

### Statistical analysis and reproducibility

Unless otherwise stated, data analyzed by parametric tests are represented by the mean ± SEM of pooled experiments and median ± interquartile range for experiments analyzed by non-parametric tests. The n-values represent the number of independent experiments performed. For each independent *in vitro* experiment, at least three technical replicates were used, and a minimum number of three experiments were performed, to ensure adequate statistical power. Student’s t-test was used to compare data with normal distribution, and non-parametric Mann–Whitney *U* test was used for samples not following a normal distribution. The confidence level used for all the statistical analyzes was of 95% (α=0.05). Two-tailed statistical analysis was applied for experimental design without predicted results, and one-tail for validation or hypothesis-driven experiments. For IC_50_ measurement, data were normalized to percent of control and the relative IC_50_ values were determined using GraphPad Prism Version 9.4.0 software. Statistical analysis was performed by GraphPad Prism software Version 9.4.0 and *R* programming language version 4.3.3.

## Results

### 13q14.2 loss is a recurrent event associated with poor breast cancer survival

Loss of chromosome cytoband 13q14.2 has emerged as one of the most frequent aberrations in patients with CLL (38). However, its frequency and clinical implications in other cancers remain largely unexplored. We initially investigated the prevalence of 13q14.2 loss across cancer types in the pan-cancer dataset from The Cancer Genome Atlas (TCGA) (39). We found that loss of this cytoband is widespread, with the highest frequencies respectively observed in non-small cell lung cancer, ovarian cancer, soft tissue sarcoma, and BCa (Fig.2).

**Fig. 2.**
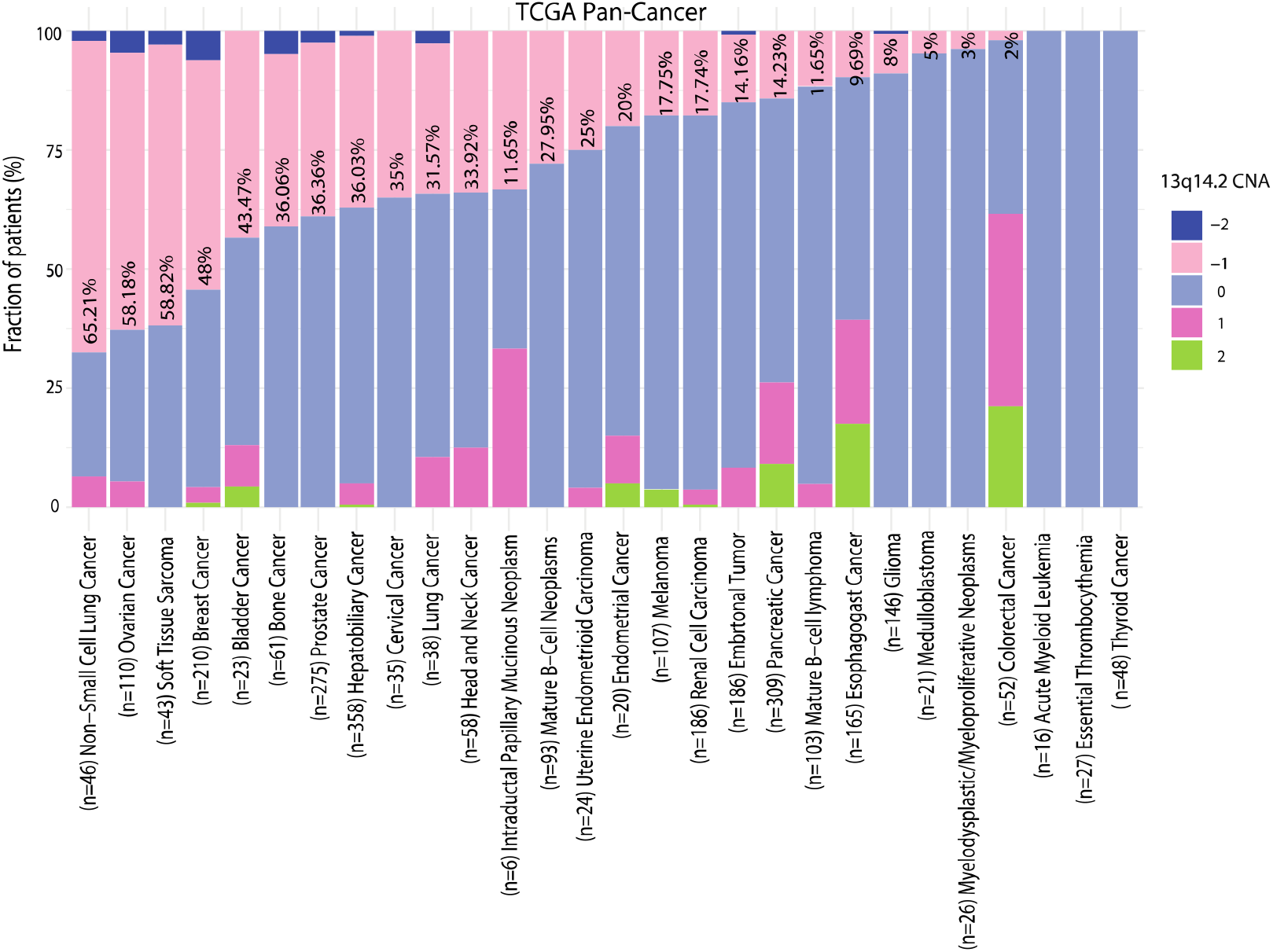
Loss of 13q14.2 is common across cancer types. Copy number alterations involving chromosome 13q14.2 were analyzed using copy number data from the TCGA pan-cancer cohort. The percentages of samples with copy number loss (−1) are shown in each bar. n: number of samples in each cancer type.-1: one copy loss,-2: two copy loss, 0: diploid copy number, +1: one copy gain, +2 two or more copy gains.

Then, given the notable occurrence of 13q14.2 loss in BCa, particularly within a substantial patient cohort (n=210), we further explored its potential role in BCa. To achieve this, we conducted a series of analyses. First, we examined the prevalence of 13q14.2 loss in comparison to other focal copy number (CN) losses using gene-based SNP6 array data in two large and well-annotated bulk BCa patient cohorts: Molecular Taxonomy of Breast Cancer International Consortium (METABRIC) (40) and TCGA. Of note, 13q14.2 loss is the 33rd most lost cytoband in the METABRIC dataset and is part of the 6th most common sub-chromosome arm-level CN loss (Fig.3a). In TCGA, 13q14.2 is the 7th most common Supplementary sub-chromosome arm-level CN (Supplementary Fig.S1a). Loss of 13q14.2 was predominantly heterozygous (−1), with 13q14.2 homozygous loss (−2) only observed in a small fraction of patients in both cohorts.

Secondly, to statistically confirm the previous data, we evaluated the frequency of 13q14.2 CNAs in comparison to other focal CNAs, encompassing both losses and gains, across the METABRIC and TCGA cohorts. Remarkably, in both datasets, the incidence of 13q14.2 loss significantly surpassed that of other losses, whereas its gain was less prevalent compared to gains observed in other regions (Fig. 3b and c). These observations indicate that the loss of 13q14.2 may be subject to positive selection pressure within tumor populations.

**Fig. 3.**
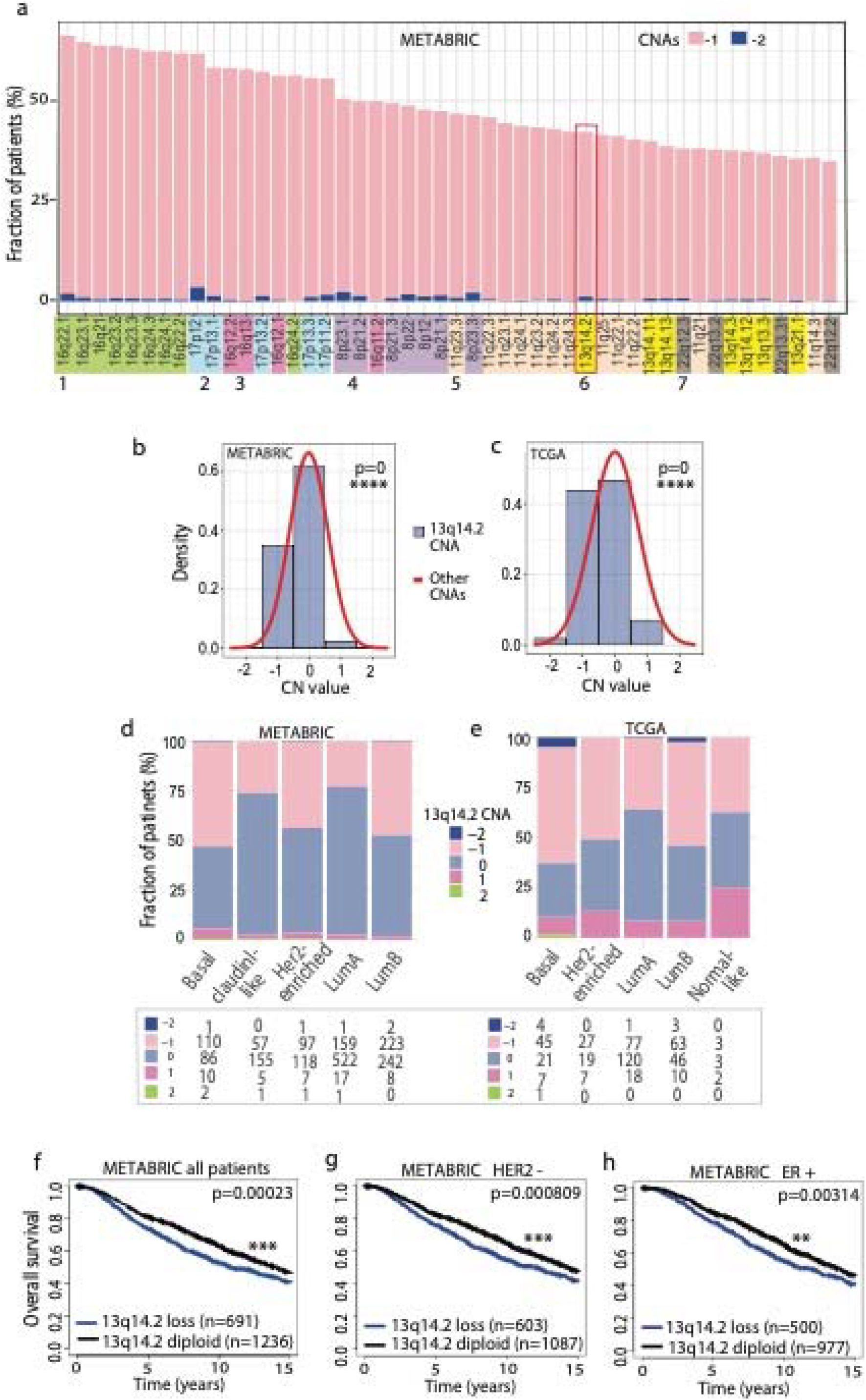
13q14.2 loss is abberent in breast cancer and is associated with poor overall survival. (a) Cytoband-level top 50 most frequent chromosomal losses, as assessed using gene-based copy number data from the METABRIC patient cohort. Cytobands on each chromosome are color-coded for easy identification. 13q14.2 is highlighted with a red box. Histograms comparing the frequencies of copy number alterations of 13q14.2 relative to other regions in the METABRIC (b) and TCGA (c) cohorts. statistics: student t-test. Relative frequencies of 13q14.2 copy number alterations assessed in the METABRIC (d) and TCGA (e) cohorts using PAM50 breast cancer classification. The numbers of patients in each subtype are shown in the box below. Kaplan-Meier plots illustrating the overall survival of breast cancer patients with 13q14.2 loss compared to those with diploid 13q14.2 within the METABRIC clinical cohort, shown for all patients (f), patients with HER2-negative (g) and ER-positive cancers (h). n: number of samples.-2: two copy loss,-1: one copy loss, 0: diploid copy number, +1: one copy gain, +2: two or more copy gains. ER: estrogen receptor. HER2: human epithelial growth factor receptor. CNA: copy number alteration. P values: log-rank test.

**Fig. 4.**
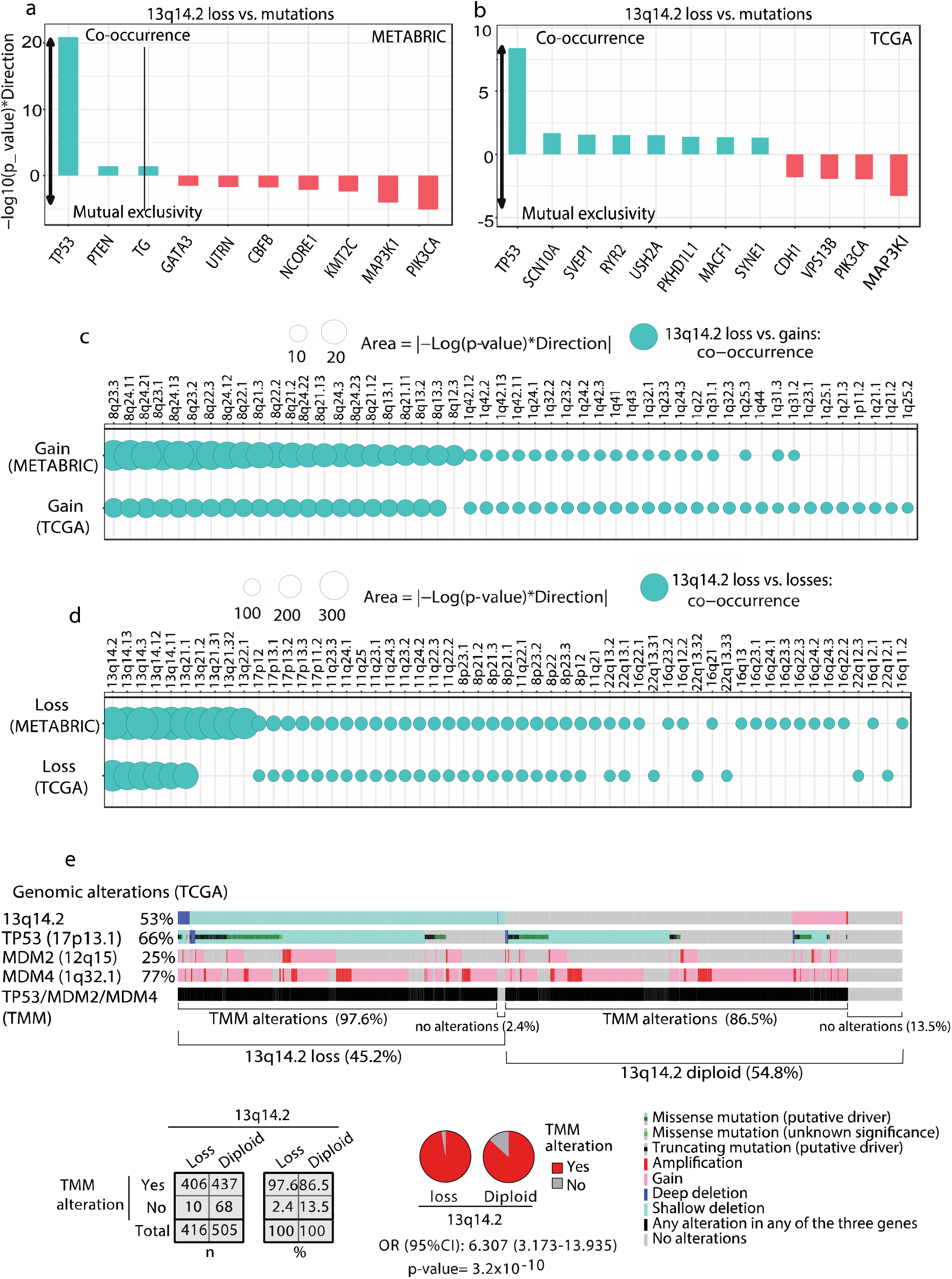
13q.14 loss accrues concurrently with other genomic alterations in breast cancers. Mutual exclusivity and co-occurrence of 13q14.2 loss with top mutated genes in (a) METABRIC and (b) TCGA breast cancer cohorts were analyzed. Mutual exclusivity and co-occurrence of 13q14.2 loss with top focal copy number alterations was analyzed in (c) METABRIC and (d) TCGA breast cancer cohorts. Statistics: Fisher’s Exact Test. (e) OncoPrint summarizing the cooccurrences of genomic alterations affecting 13q14.2, TP53, MDM2 and MDM4 in the TCGA BCa cohort. Statistics: Fisher’s exact test with odds ratio (OR) and 95% confidence interval (CI).

**Fig. 5.**
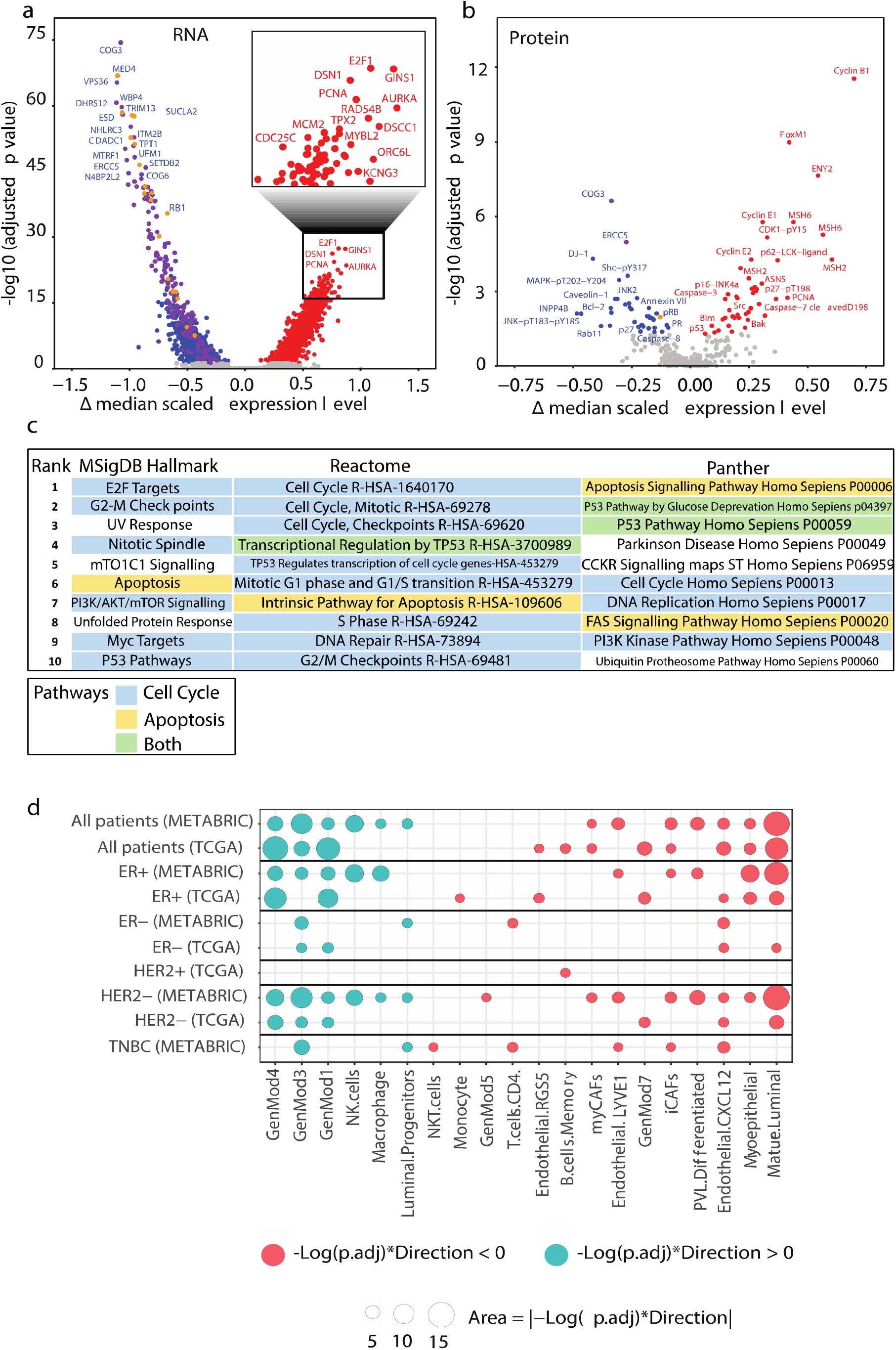
13q14.2 loss activates cell cycle and apoptosis-related pathways in cancer cells and affects the tumor microenvironment. (a) Volcano plot demonstrating differential gene expression analysis in TCGA patient samples with and without 13q14.2 loss. Blue dots: genes significantly downregulated. Red dots: genes significantly upregulated (i.e., adjusted p value<0.05). Orange circles: genes on the 13q14.2. Purple circles: all other genes on chromosome arm 13q. Red circles: upregulated genes. (b) Volcano plot demonstrating differential (phospho-) protein expression analysis in TCGA patient samples with and without 13q14.2 loss. Blue dots: proteins significantly downregulated. Red dots: proteins significantly upregulated (i.e., adjusted p value<0.05). Orange circles: proteins whose genes reside on 13q14.2. Purple circles: all other proteins whose genes reside on chromosome arm 13q. (c) Pathway analyses performed on the upregulated genes and proteins in TCGA samples with 13q14.2 loss using three different pathway enrichment tools, including MSigDB hallmarks, Reactome and Panther. (d) Comparison between fractions of cell types in tumors with 13q14.2 loss versus 13q14.2 diploid in METABRIC and TCGA cohorts.

Thirdly, we explored the frequency of 13q14.2 CNAs within distinct BCa subtypes in METABRIC and TCGA, delineated by the Prediction Analysis of Micro-array 50 classification (PAM50) (41). Notably, approximately 50% of patients classified as Basal, Her2-enriched, and lumB subtypes exhibited 13q14.2 heterozygote loss, while about 25-30% of patients identified as LumA and Claudin-like subtypes, manifested this loss in both cohorts (Fig. 3d and e). Less than 2% of all patients exhibited amplification of 13q14.2 in these cohorts.

Lastly, we utilized the clinical data from the METABRIC and TCGA datasets to explore the impact of 13q14.2 loss on patient overall survival. Patients with 13q14.2 loss demonstrated markedly poorer survival outcomes compared to their wildtype counterparts within the METABRIC cohort (Fig.3f). Further stratification by molecular subtypes unveiled significant survival discrepancies, particularly within the HER2-and ER+ subgroups (Fig.3d and e). Although statistical significance was not reached, a trend toward poorer survival was observed for patients harboring 13q14.2 loss compared to those with 13q14.2 diploidy in other subtypes of the METABRIC cohort (Supplementary Fig. S1b-d) and across the TCGA cohort (Supplementary Fig. S1e-i).

These collective findings underscore the prevalent occurrence of 13q14.2 loss in BCas, its association with adverse survival outcomes in specific molecular subtypes, and its potential as a clinically relevant biomarker.

### 13q14.2 loss in breast cancer is part of a larger repertoire of genomic alterations rather than an isolated event

In cancer cells, multiple evolutionary pathways are pursued to pro-mote survival and growth. Oncogenic drivers compete under selective pressure, leading to the dominance of certain drivers while others are outcompeted, resulting in mutual exclusivity (42, 43). However, certain oncogenic drivers may also be selected concomitantly, most likely due to a co-regulatory function during tumorigenesis (43). To ascertain whether 13q14.2 loss represents an independent oncogenic event or if it coincides with other genomic alterations in BCas, we performed co-occurrence and mutual exclusivity analyses with both mutated genes and other recurrent CNAs in TCGA and METABRIC BCa cohorts. Firstly, we conducted an analysis of mutual exclusivity or co-occurrence using Fisher’s exact test, which involved computing *p*-values and odds ratios for each gene among those with a mutation frequency greater than 2% against the loss of 13q14.2 in both the TCGA and METABRIC cohorts. Mutated genes exhibiting a positive-log10(*p* value) * Direction, such as *TP53*, strongly co-occurred with 13q14.2 loss (Fig.4a and b). The observed co-occurrence of *TP53* mutations with loss of 13q14.2 is particularly noteworthy, given that *TP53* serves as the gatekeeper of the cell cycle (44). Loss of the 13q14.2 cytoband, which includes the *RB1* gene that controls the cell cycle, combined with *TP53* mutation, synergistically leads to a significant reduction in the cell’s ability to regulate its growth and repair DNA damage. This identifies a potential treatment target for tumors harboring loss of the 13q14.2 cytoband. Conversely, mutated genes exhibiting negative-log10(p value) * Direction, including *MAP3K1* and *PIK3CA*, were mutually exclusive with 13q14.2 loss in both cohorts (Fig.4a and b). This suggests that the regulatory effect of 13q14.2 loss might represent both alternative or overlapping pathways for tumor development with that of involving mutations in *MAP3K1* and *PIK3CA*.

Secondly, we conducted a similar analysis to assess the co-occurrence or mutual exclusivity of 13q14.2 loss with the loss or gain of other top altered cytobands in both the TCGA and METABRIC cohorts. Notably, loss of 13q14.2 often coincides with gains of 8q12-q24 and 1q22-43 (Fig.4c), as well as losses of cytobands adjacent to 13q14.2 and 17p11-p13, 11q22-q25, and 8p12-p23 (Fig.4d) across both patient cohorts. It is noteworthy that the tumor suppressor gene *TP53* (17p13.1) and its negative regulator *MDM4* (1q32.1) are located within the co-occurring loci.

To further explore this, we conducted additional analyses of the genomic alterations involving these genes in TCGA BCa samples, comparing those with and without 13q14.2 loss. This revealed that *TP53*/*MDM2*/*MDM4* alterations occurred 11% more frequently in cases where 13q14.2 was lost (Fig.4e; p=3.2×10-10, Fisher’s exact test). This suggests a potential functional relationship between the loss of 13q14.2 and *TP53/MDM2/MDM4*, involving both mutations and CNAs. The co-occurrence of these alterations may indicate a synergistic effect or shared underlying mechanisms in BCa development or progression.

These observations collectively suggest that 13q14.2 loss does not act alone but interacts with other genomic alterations to drive tumor development.

### 13q14.2 loss affects breast cancer cells and their tumor microenvironment

To study the impact of 13q14.2 loss on cancer cells, we conducted differential gene expression analysis (DEA) using mRNA data from patient samples with 13q14.2 loss compared to those with diploidy for this region, utilizing the TCGA BCa dataset. As expected, the expression of multiple genes located in the 13q14.2 region were significantly downregulated (Fig.5a, orange circles). Interestingly, other genes located on the 13q chromosome arm were also downregulated, suggest-ing that a considerable number of patients have lost larger fractions of the 13q chromosome arm in addition to 13q14.2 loss (Fig.5a, purple circles). In contrast, multiple genes involved in cell cycle-related pathways were remarkably upregulated, including but not limited to *E2F1*, *DSN1*, *GINS1*, *PCNA*, *AURKA*, *TPX2* and *DSCC1* (Fig.5a, red circles).

To complement this gene expression analysis, we also performed differential protein expression analyses in the same patient samples, where available. It is worth noting that the number of statistically significant downregulated (phospho-) proteins on 13q14.2 (n=1) and 13q (n=3 including the one on 13q14.2) was considerably lower than at the mRNA level. This difference can be attributed to the fact that fewer (phospho-)proteins were analyzed, and post-transcriptional compensatory mechanisms that may affect protein levels. Consistent with the gene expression data, protein levels of many cell cycle-related proteins were significantly increased, including *CyclinB1*, *FoxM1*, and *CyclinE1*. Additionally, we observed upregulation of several pro-apoptotic components, including *Caspase-7-cleaved D198*, *BAK*, *BIM*, and *Caspase-3* (Fig.5b, red circles). Conversely, anti-apoptotic proteins like *BCL2* were downregulated (Fig.5b).

These observations prompted us to perform pathway analysis using the combined mRNA and protein differential expression data. To avoid potential biases introduced by downregulated genes on 13q14.2 due to CN loss that may not reflect true functional differences, we focused on upregulated genes and proteins. By analyzing the top significantly upregulated mRNAs and proteins, using sources including ‘MSigDB Hallmark’(45) ‘Reactome’(46) and ‘Panther’(47), we identified two prominent upregulated pathways: ‘Cell cycle’ and ‘Apoptosis’ (Fig.5c). Interestingly, the loss of the 13q14.2 region is associated with up-regulation of the *PI3K/AKT/mTOR* and *p53* pathways.

To explore the potential impact of 13q14.2 loss on the cellular composition in the tumor microenvironment, we estimated the fractions of different cell types within the breast tumor microenvironment. Employing our established pipeline (23), we utilized high-resolution scRNA-seq data (24) as a reference for the breast TME and employed CIBERSORTx as a deconvolution method (22). The scRNA-seq data encompassed a diverse array of cell types, including cancer cells divided into seven recurrent gene modules (GenMod1-7). Each module is enriched in specific biological pathways, such as *Jun*, *Fos*, *ER*, *RTK*s, and *TP53* for GenMod1; *Myc* and *OxPhos* for GenMod2; and EMT, IFN, and Complement for GenMod3, among others (24). Our analysis revealed that loss of 13q14.2 resulted in a higher fraction of GenMod4, characterized by proliferating cancer cells (Fig.5d). This finding aligns with our DEA, which indicated that loss of 13q14.2 led to enrichment of cell cycle-related proteins. Additionally, we observed a significant decrease in the fractions of healthy epithelial and endothelial cells, such as mature luminal cells and myoepithelial cells, associated with 13q loss, consistent with the presence of fewer differentiated cells (Fig.5d). Furthermore, 13q14.2 loss led to a higher fraction of natural killer (NK) cells and macrophages in METABRIC cohort but not in the TCGA cohort (Fig.5d). Analysis of the cell fractions enumerated from spatial omics data in the METABRIC cohort confirmed a positive association between 13q14.2 loss and an increased number of macrophages (Supplementary Fig.S2).

These findings collectively suggest that 13q14.2 loss contributes to alterations in the composition of the tumor microenvironment, favoring the proliferation of cancer cells.

**Loss of 13q14.2 sensitizes breast cancer cells to *BCL2* inhibitors.** Following a Sanger Institute pipeline (26, 48), we previously applied a machine learning approach to a pan-cancer drug screen dataset to identify mutations and focal or chromo-some arm-level CNAs associated with increased drug sensitivity or resistance (27). Here, we leveraged an expanded version of this dataset to apply this approach on a breast cancer-specific subset of these data, including 30,394 half-maximum inhibitory concentration (IC_50_) values from 542 unique drugs across 52 BCa cell lines (28). Notably, using this unbiased approach, we identified a novel CNA-drug association with robust impact: loss of 13q14.2 strongly associates with increased sensitivity of BCa cells to *BCL2*-targeting drugs (Fig.6a, blue circles). This association was also identified by the Sanger Institute analysis (Fig.6a, orange triangle) using a prior version of the dataset, albeit at a lower statistical significance level (27).

We note that our analysis also identified genomic amplification of the *HER2* gene, located on chromosome 17q12, as strongly associated with sensitivity to *HER2-* targeting drugs in BCa cells (Fig.6a, yellow squares). Identification of this well-established interaction (49) thus validating our approach.

To delve deeper into the association between 13q14.2 loss and *BCL2* inhibitors, we investigated which drugs exhibited the most substantial effect sizes and increased sensitivity to 13q14.2 loss in the BCa cell lines across 673 tested drugs. Intriguingly, *BCL2* inhibitors, including venetoclax, ABT737, and navitoclax, dis-played the most significant associations with increased sensitivity in BCa cell lines with 13q14.2 loss (Fig. 6b, green circles). However, we noted that 13q14.2 loss did not statistically significantly associate with other *BCL2* inhibitors. In addition, 13q14.2 loss did not statistically significantly associate with resistance to any of the 673 drugs (Fig. 6b).

**Fig. 6.**
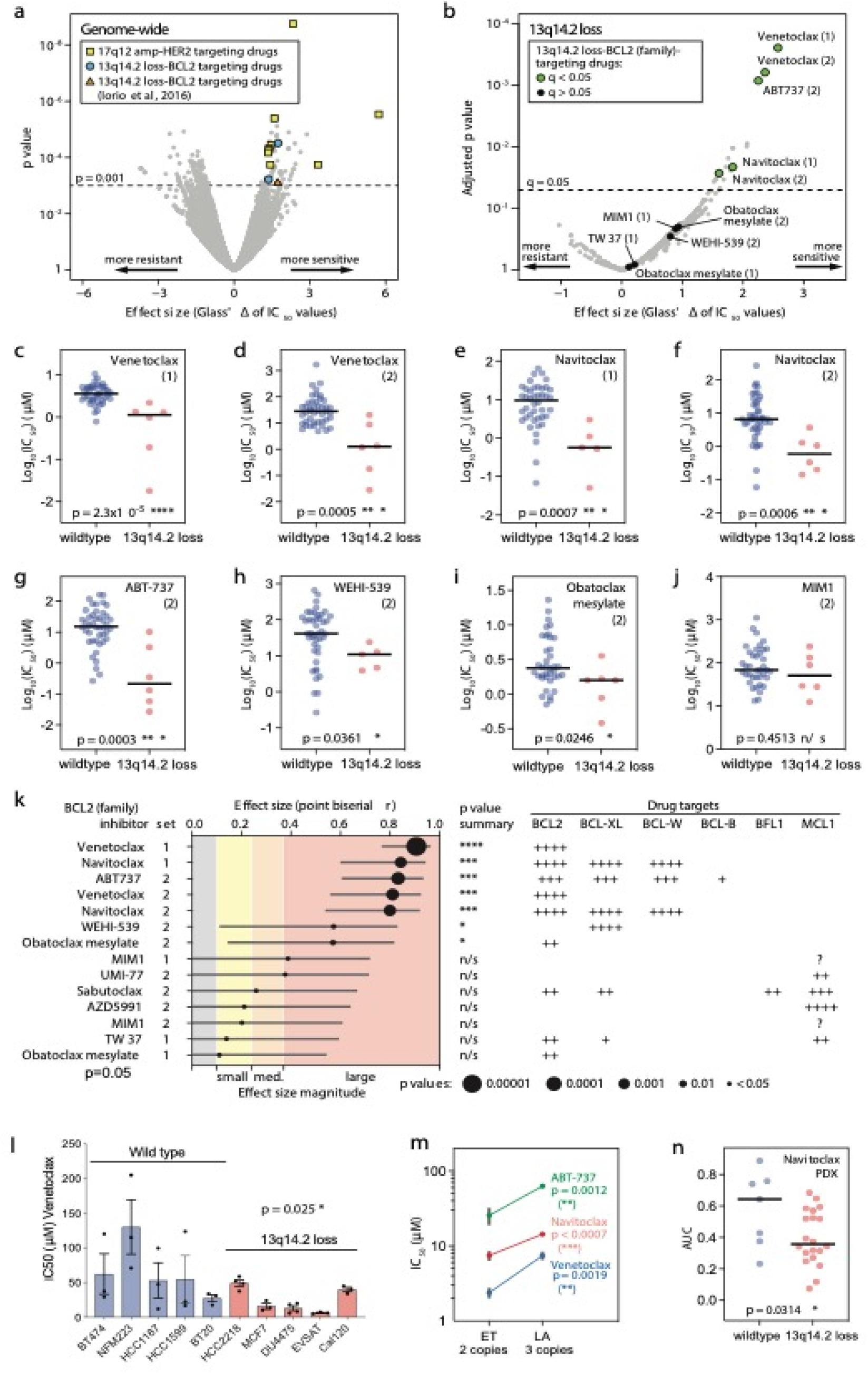
13q14.2 loss is associated with sensitivity to *BCL2* inhibitors in breast cancer. (a) Volcano plot showing the results of an elastic modelling to evaluate the association of 410 established genomic features (high confidence cancer gene mutations and recurrent focal CNAs) with IC50 values of 438 drugs across 52 breast cancer cell lines. Selected established (involving HER2) and new pharmacogenomic interactions (involving 13q14.2 loss) are highlighted. Cut-offs: p value: 0.001, FDR:0.25, Glass’ Δ effect size: 1.4. (b) Volcano plot evaluating the associations between 13q14.2 loss and all drugs in a panel of 673 tested drugs from two independent screens (1 and 2, respectively), including some drugs tested in both screens. The number of unique drugs is smaller. Cut-offs: adjusted p value (q value): 0.05, FDR:0.25, Glass’Δ effect size:1.4. Pharmacogenomic interactions involving *BCL2* inhibitors are highlighted in green (q<0.05) and black (q<0.05). (c-j) Scatter dot plots comparing the log10 (IC50) of BCL2 inhibitors from the two screens in BCas cell lines with wild type 13q14.2 (blue) and 13q14.2 loss (red). P values: Two-tailed Mann-Whitney U tests. * p value<0.05, ** p value<0.01, *** p value<0.001, **** p value<0.0001. (k) Left: Effect size magnitudes for each of the *BCL2* inhibitors are shown in different colors across all BCa cell lines possessing 13q14.2 loss. The circle sizes correspond the p values. This analysis includes duplicates, i.e., same drugs tested in both screens, included in dataset 1 (set=1) and dataset 2 (set=2). Right: Target proteins of individual drugs are shown. Magnitudes of inhibitory effects are shown by increasing numbers of ‘+’ symbols; ?: unknown effect. (l) IC50 sensitivity to venetoclax was assessed in 5 breast cancer cell lines with wild type 13q14.2 and 5 breast cancer cell lines with 13q14.2 loss. Statistics: one-tailed Mann-Whitney U test. (m) IC50 sensitivity to venetoclax, navitoclax and ABT-737 was assessed in a breast cancer cell line with two copies of chromosome 13 (PMC42-ET) and a PMC42-ET-derived breast cancer cell lines with an extra copy of chromosome 13 (PMC42-LA). Vertical grey bars show the range of IC50 values determined in three independent replicate experiments with the median values shown as dots. P values: Paired t tests. (n) Dot plot graph comparing the response to navitoclax between breast cancer patient-derived xenografts with 13q14.2 loss and their wildtype counterparts. Data extracted from (Bruna et al., 2016). AUC: area under the ROC curve. P value: Two-tailed Mann-Whitney U test.

Classifying the BCa cell lines into two groups, those with loss of 13q14.2 and those wild type (diploid) for this region, we then compared their sensitivities to these 7 *BCL2* inhibitors. This showed that loss of 13q14.2 resulted in significantly lower IC_50_ values (indicating greater sensitivity) for venetoclax, navitoclax, ABT-737 and WEHI-539, including in two independent screens for the former two (Fig.6c-i), but not for and MIM1, sabutoclax, TW37, AZD5991 and UMI-77 (Supplementary Fig.S3a-f). We observed a marginally increased sensitivity for obatoclax in one screen (Fig.6i) but not in another screen (Supplementary Fig.S3c).

To further characterize the sensitivities of BCa cell lines with 13q14.2 loss to *BCL2* inhibitors, we determined the effect size magnitude of each inhibitor. Venetoclax exhibited the largest effect size with the lowest *p* value <0.00001, followed by navitoclax and ABT-737 with *p* value <0.0001. WEHI-539 and obatoclax ranked third with *p*values <0.01 and < 0.05, respectively (Fig.6k, left). Notably, comparison of the selective inhibitory ef-fects of each drug towards different *BCL2* family member pro-teins strongly suggests that these increased sensitivities are primarily, if not exclusively, due to inhibition of *BCL2* protein, rather than other *BCL2* family member proteins (Fig.6k-right).

To validate the increased sensitivity of BCa cells with 13q14.2 loss to *BCL2* inhibitors *in vitro*, we treated a panel of 10 BCa cell lines with and without 13q14.2 loss venetoclax, the most specific inhibitor of *BCL2*. The presence or absence of 13q14.2 loss and its approximate size within the genome was initially confirmed by SNP6 microarray (Supplementary Fig.S4b-h) and shallow sequencing (Supplementary Fig.S5). Subsequently, the cell lines were treated with serial log*10*-fold concentration dilutions of venetoclax, and their viability was measured after 72 hours. Importantly, cells with 13q14.2 loss exhibited significant sensitivity to venetoclax compared to their wildtype counterparts (Fig.6l and Supplementary FigS3g-p) (One-tailed Mann-Whitney u-test, p value= 0.025). It is worth noting that the response to venetoclax varied across the cell lines harboring a loss of 13q14.2 with EVSA-2 being the most sensitive and HCC2218 the least sensitive to venetoclax (Fig.6l). The variation was also seen in the cell lines with diploid or amplified 13q14.2 with MFM223 the most resistant and BT20 the least resistant cell lines to venetoclax (Fig.6l).

To evaluate whether the presence or absence of *BCL2* protein affects the association between loss of the 13q14.2 region and sensitivity to venetoclax in cell lines, we measured the protein levels of *BCL2*. However, our data suggest that the relationship between 13q14.2 deletion and venetoclax sensitivity appears to be independent of *BCL2* expression (Supplementary Fig.S3q).

To investigate whether amplification of this region influences the response to *BCL2* inhibitors, we employed two BCa cell lines: PMC42-ET, possessing two copies of chromosome 13, and PMC42-LA, derived from PMC42-ET with three copies of chromosome 13 (37) (Supplementary Fig.S6a-b). We evaluated the chemotherapeutic sensitivities of PMC42-ET and PMC42-LA to Venetoclax (ABT-199) and two pan-specific *Bcl-2/Bcl-XL* inhibitors, ABT737 and Navitoclax (ABT263). The IC_50_ values for each drug were determined by treating the cell lines with serial 10-fold dilutions of the drugs and measuring their viability after 72 hours. Our drug assays consistently showed that PMC42-LA cells, with an extra copy of chromosome 13, exhibited significantly higher resistance to all three drugs compared to PMC42-ET (Fig.6m). The IC_50_ values for ABT199 were 2.60 µM and 9.12 µM for parental PMC42-ET and derivative PMC42-LA, respectively; for ABT737, the values were 0.83 µM and 1.23 µM; and for ABT263, the values were 7.46 µM and 14.36 µM (Supplementary Fig.S6c-e). Consequently, our observation that cells harboring an extra copy of chromosome 13 exhibit increased resistance to three *BCL2* inhibitors aligns with our findings that the loss of the 13q14.2 region enhances the sensitivity of BCa cell lines to these drugs.

Considering that patient-derived xenograft models more faith-fully replicate the molecular characteristics and tumor environment of patients, we expanded our analysis from cell lines to data obtained from a biobank of BCa patient-derived xenograft explants (50). We aimed to determine the effect of 13q14.2 loss on the sensitivity of patient-derived xenografts (PDXs) to navitoclax compared to their wildtype counterparts. Notably, the in-creased sensitivity to the drug resulting from 13q14.2 loss was also evident in these PDX models (Fig.6n), providing further support for the significant role of this CNA in the treatment response to *BCL2* inhibitors.

These data collectively suggest that loss of 13q14.2 enhances the therapeutic response to *BCL2* inhibitors. Consequently, loss of 13q14.2 could potentially serve as a predictive biomarker for the efficacy of *BCL2* inhibitors in treating BCa.

## Discussion

While *RB1* on the 13q14.2 locus has been extensively studied due to its established tumor suppressor function (51–53), the 13q14.2 region encompasses additional genes that may significantly in-fluence the development and progression of BCa beyond *RB1*. This perspective is supported by both publicly available sequencing data and our own research, which demonstrate the variable extent of 13q14.2 losses across various BCa cell lines and patient samples (Supplementary Fig.S7). The impact of other potential genes in this region is particularly underscored by studies identifying the concurrent loss of *RB1* and *lysophosphatidic Acid Receptor 6 (LPAR6)* at the 13q14.2 locus, a combination that has been shown to increase the proliferation rate of BCa cells (51, 54). Moreover, the relationship between larger CNAs and increased disease aggressiveness, along with their significant implications for clinical outcomes, has lately been a subject of intensive research focus (40, 55, 56). In hematological cancers such as CLL, loss of chromosome 13q14 is a predominant event, causing poor survival (11). Our findings revealed that in solid tumors, recurrent loss of 13q14.2 is a defining feature of the genomic landscape, underscoring its significance in tumor pathogenesis. We focused on the prominence of chromosome 13q14.2 loss as a prevalent chromosomal aberration across BCa subtypes, occurring in approximately one-third of all BCas cases from the TCGA and METABRIC cohorts. The significant selective pressure favoring the loss of one copy of chromosome 13q14.2 over other CNAs suggests a fitness advantage for tumor cells, ultimately probably contributing to poorer overall survival outcomes for patients.

Recent findings highlight a significant mutual exclusivity between common mutations and CNAs in cancers. For example, in pancreatic cancer, *kirsten rat sarcoma viral oncogene homologue (KRAS)* mutations are mutually exclusive with chromosome 18q gains, both targeting overlapping signaling pathways (57). Similarly, in our BCa study, mutual exclusivity between *PIK3CA* and *MAP3K1* mutations and the loss of 13q14.2 raises intriguing questions about the functional redundancy or compensatory mechanisms within cancer cells. *PIK3CA*, encoding a subunit of PI3K, directly influences the PI3K/AKT pathway, a critical route for promoting oncogenic activity including cell growth and survival. In our differential expression analyses, we identified activation of the PI3K/AKT pathway in response to 13q14.2 loss. This suggests that the loss of 13q14.2, potentially through the deletion of inhibitors of the PI3K/AKT pathway or through indirect effects on pathway regulators, mimics the activating effects of *PIK3CA* mutations. Such redundancy underscores the cell’s ability to maintain crucial oncogenic signaling by alternative genomic routes, which may compensate for the absence of mutations in key pathway genes like *PIK3CA per se*.

The co-occurrence of both *TP53* CN loss and mutations with 13q14.2 loss in BCa samples indicates a complex regulatory mechanism involving both alleles of *TP53*. In many cancers, *TP53* function is disrupted. This may occur via a range of mechanisms, including *TP53* mutation, *TP53* CN loss, or gain or amplification of genes encoding *TP53* antagonizing proteins, like *MDM2* and *MDM4* (58, 59). For instance, gain of chromosome 1q, which harbors the *MDM4* gene and occurs in many BCas, was recently shown to be largely mutually exclusive with *TP53* mutations, but associated with downregulation of *TP53* target genes (60). Moreover, a recent study has shown that *TP53* LOH results in genomic instability, which leads to chromosome 13 loss, along with other CNAs, during the progression of pancreatic ductal adenocarcinoma (61). These notions are consistent with our observation that 13q14.2 loss statistically significantly co-occurs with *TP53*-inactivating genomic events and support our hypothesis that the loss of 13q14.2 in BCa could be a downstream consequence of initial *TP53* pathway-impairing genomic alterations.

CNAs as a ubiquitous feature of cancers have been attributed to the imbalance in transcriptomic and proteomic landscapes of tumor cells (62, 63). We observed a significant upregulation of several genes and proteins associated with cell cycle and pro-apoptotic pathways, as well as downregulation of anti-apoptotic genes, such as *BCL2*, in patient samples harboring 13q14.2 loss. We envisage two possible scenarios for these observations: 1. Apoptosis, a critical cell death program, plays a dual role in cancer. While primarily known for its tumor-suppressing effects, apoptosis also has pro-oncogenic properties (64). The pro-oncogenic property is mainly due to its non-cell-autonomous effects, particularly involving the interaction with tumor-associated macrophages in the tumor microenvironment (61, 65, 66). This correlates with our findings that demonstrate the positive impact of 13q14.2 loss on the macrophage populations within the TME. When apoptotic tumor cells are cleared by macrophages —a process known as efferocytosis— it can lead to an anti-inflammatory and pro-tumorigenic environment (67). This process often results in the secretion of cytokines and growth factors that can further promote tumor growth and metastasis. 2. As demonstrated in our study, additional genomic or pathway alterations, such as alterations in *TP53* or activation of the *PI3K/AKT* pathway — both known for inhibiting apoptosis and promoting cell survival — often coincide with 13q14.2 loss. Of note, these alterations can counteract the effects of activated apoptotic pathways, enabling cancer cells not just to survive but to thrive, enhancing their malignant potential (68, 69).

Our data suggest that 13q14.2 loss in BCas may confer a selective advantage by promoting oncogenic behaviors, enhancing tumor growth and survival. Yet, intriguingly, this same genomic alteration also introduces vulnerabilities that can be exploited for therapeutic purposes. Our research has revealed that the loss of 13q14.2 sensitizes BCa cells to *BCL2* inhibitors, offering a unique, paradoxical opportunity for targeted intervention. This dual impact of 13q14.2 loss highlights the complex, interwoven regulatory networks within cancer cells, where a single genomic alteration can both bolster cancer cell survival and introduce critical susceptibilities. This phenomenon aligns with the concept of synthetic lethality, where the interaction between two genomic alterations — each tolerable on their own — can cumulatively lead to cell death (70). Specifically, the combination of 13q14.2 loss and inhibition of *BCL2* exemplifies synthetic lethality, presenting a double strike that can significantly hinder cancer cell viability and offer a strategic target for BCa therapy. It is worth noting that the *BCL2* inhibitor venetoclax is recognized for its dual mechanism of action: it not only inhibits the *BCL2* protein but also induces metabolic rewiring independently of *BCL2* inhibition (71). In our studies, we have observed a *BCL2*-independent response to venetoclax in BCa cell lines, along with a variable response in cell lines harboring 13q14.2 loss. This suggests that partial or extensive losses within this region, which involve a varying number of genes, might play a role in modulating drug sensitivity.

While our study highlights the relevance of 13q14.2 loss in BCa, it also uncovers a significant gap. Notably, there is a lack of an isogenic *in vitro* model that specifically targets 13q14.2 loss. This is crucial for excluding cell-autonomous effects of other genomic alterations present in cell lines and for a detailed mechanistic study of the impact of 13q14.2 loss on BCa cell biology and drug response. The recent adoption of CRISPR-CAS9 technology to facilitate the development of a stable cell line with specific CNAs (60, 72, 73) can enable more precise investigations into the biological consequences of 13q14.2 loss.

## Conclusion

Our research into the loss of chromosome 13q14.2 in BCa reveals its profound impact on both the genomic and therapeutic landscapes of the disease. This chromosomal aberration is associated with poor patient survival, likely by promoting oncogenic traits, such as anti-apoptotic and proliferative phenotypes. However, it also introduces specific vulnerabilities that can be exploited using *BCL2* inhibitors, showcasing the potential of synthetic lethality as a strategy in cancer therapy. This dual role highlights the importance of understanding genomic alterations for developing targeted cancer treatments.

## List of abbreviations

BCa: Breast Cancer
CNA: Copy Number Alterations
CIN: Chromosomal Instability
CLL: Chronic Lymphocytic Leukemia
CRISPR-CAS9: Clustered Regularly Interspaced Short Palindromic Repeats and CRISPR-associated protein 9
DAE: Differential Expression Analysis
LOH: Loss Of Heterozygosity
METABRIC: Molecular Taxonomy of Breast Cancer International Consortium
PDX: Patinet Derived Xenograft
PAM50: Prediction Analysis of Microarray 50
TCGA: The Cancer Genome Atlas
TME: Tumor Microenvironment

## Declarations

### Ethics approval and consent to participate

Not applicable

### Consent for publication

Not applicable

### Availability of data and material

All R codes required to replicate the analysis are available at https://github.com/Paris-Shahrouzi/Parastoo-Shahrouzi-13q14.2-loss.git. All data generated or analysed during this study are included in this published article [and its supplementary information files]. The datasets supporting the conclusions of this article are available in the [TCGA PanCancer Atlas study and BRCA METABRIC (2016)], [https://www.cbioportal.org]. For bioinformatic analysis R programming language version 4.4.0 was emplyed. Geraphical abstract was created with BioRender.com.

## Authors’ contributions

P.S. wrote this manuscript, performed the majority of bioinformatic and *in vitro* analysis and designed some of the experiments. Y.A has performed several bioinformatic analysis. W.B has supported the *in vitro* experiments. S.B has performed the *in vitro* validation of the effect of 13q amplification on drug response. D.K has supported the *in vitro* analysis. D.R provided support during the execution of experiments in the translational research institute in Brisbane, Australia. X.T supported the bioinformatic analysis. V.N.K and P.H.G.D. conceived the study and supervised the execution of the project. All authors have read and agreed to the published version of the manuscript.

## Acknowledgment

We thank Dr. Bauke Ylstra and Dr. Erik Van Dijk for performing and analyzing the shallow whole genome analysis. We thank Dr. Farzaneh Forouz for helping with the analysis of SNP6 data using GenomeStudio software.

