## Supplementary data for "Loss of chromosome cytoband 13q14.2 orchestrates breast cancer pathogenesis and drug response"

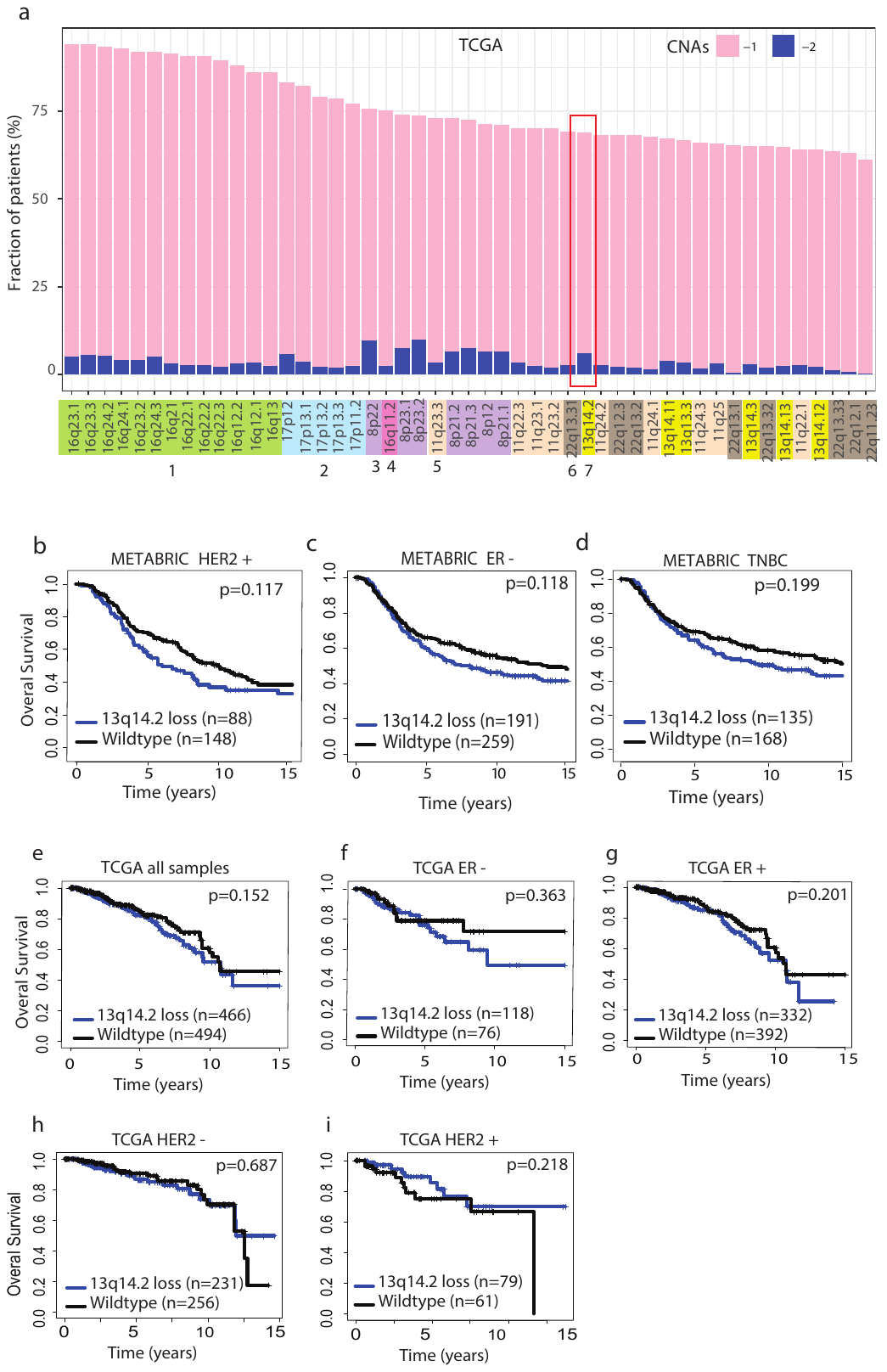

**Fig. S1. (a)** The chromosomal loss or deletion of top 50 cytobands were plotted from TCGA GISTIC copy number data. 13q14.2 is highlighted with a red box -1: one copy loss. -2: two copy loss. Kaplan Meier assessing the overall survival associated with 13q14.2 loss in comparison with the wildtype counterparts was performed in the METABRIC **(b)** HER2 negative, **(c)** ER negative and **(d)** TNBC samples. Kaplan Meier assessing the overall survival associated with 13q14.2 loss in comparison with the wildtype counterparts was performed in TCGA **(e)** all samples, **(f)** ER-negative, **(g)** ER-positive, **(h)** HER2-negative and **(i)** HER2-positive samples. n: number of samples, ER: estrogen receptor, TNBC: triple negative breast cancer, HER2: human epithelial growth factor receptor. log rank test p value is denoted.

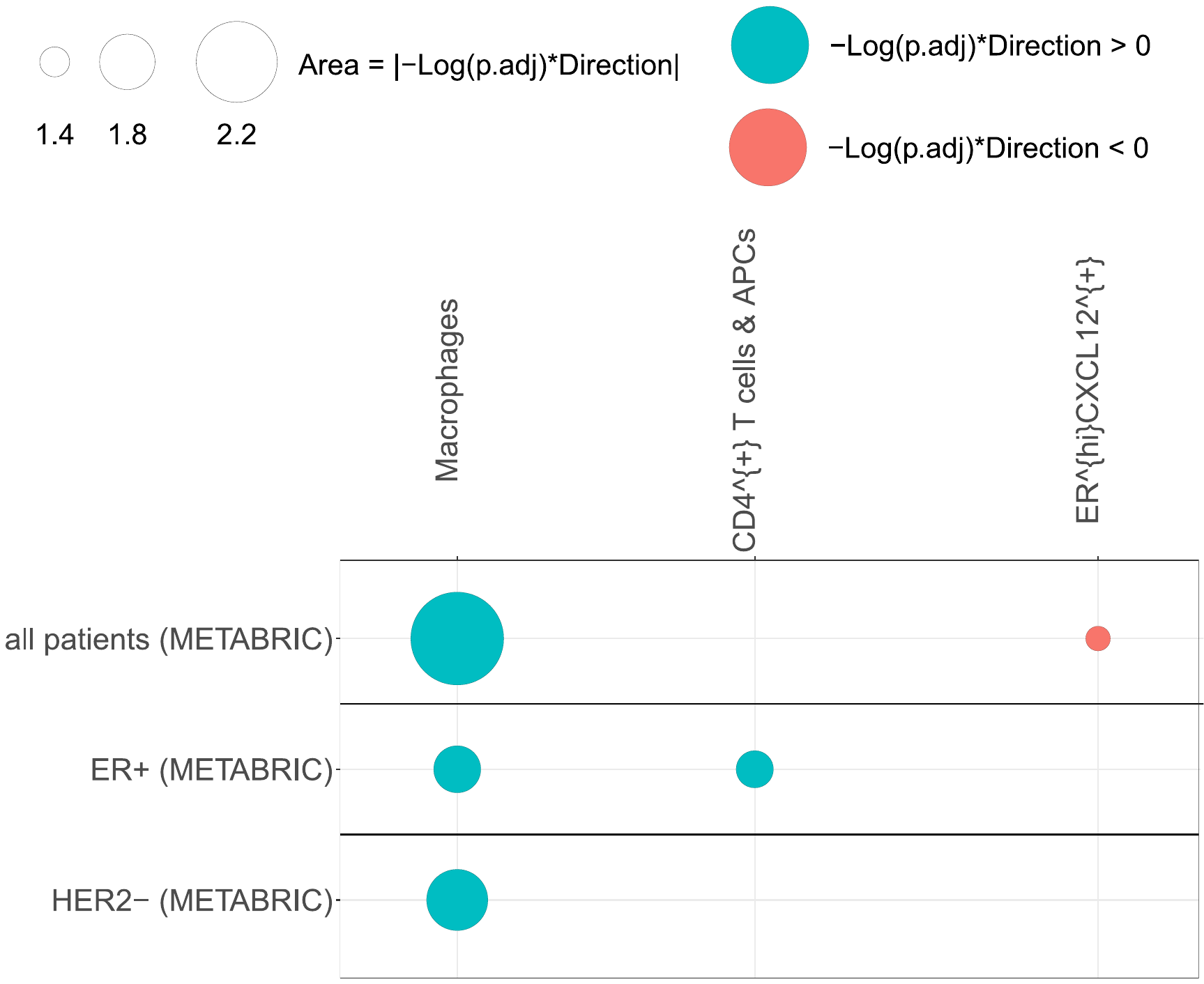

**Fig. S2.** Comparison of cell fractions enumerated from spatial omics data in tumors with 13q loss vs wild type. Imaging mass cytometry data was employed for a subset of METABRIC cohort (25) which were downloaded from <https://zenodo.org/record/5850952> and fraction of different cell types were compared.

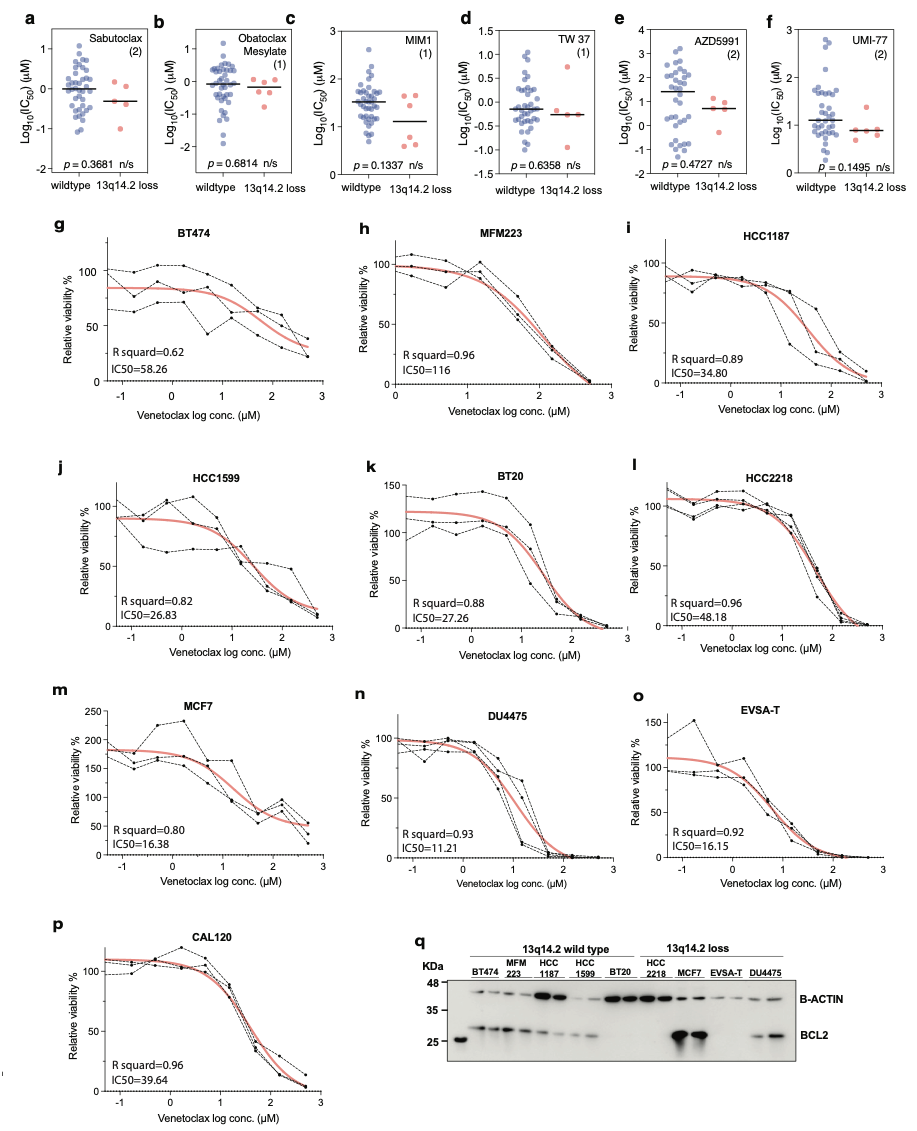

**Fig. S3.** The association of 13q14.2 loss with the sensitivity to BCL2 inhibitors. **(a-f)** the comparison of the log10 (IC50) of BCL2 inhibitors from two screens in BCas cell lines with wild type 13q14.2 (blue) and 13q14.2 loss (red). This includes duplicates, i.e., same drug tested in both screens; hence for example ‘AZD5991 (2)’ refers to 2 different screens. **(g-p)** IC50 measurement of log2 concentrations of venetoclax in the indicated breast cancer cell lines, measured by GlowTiter viability assay.

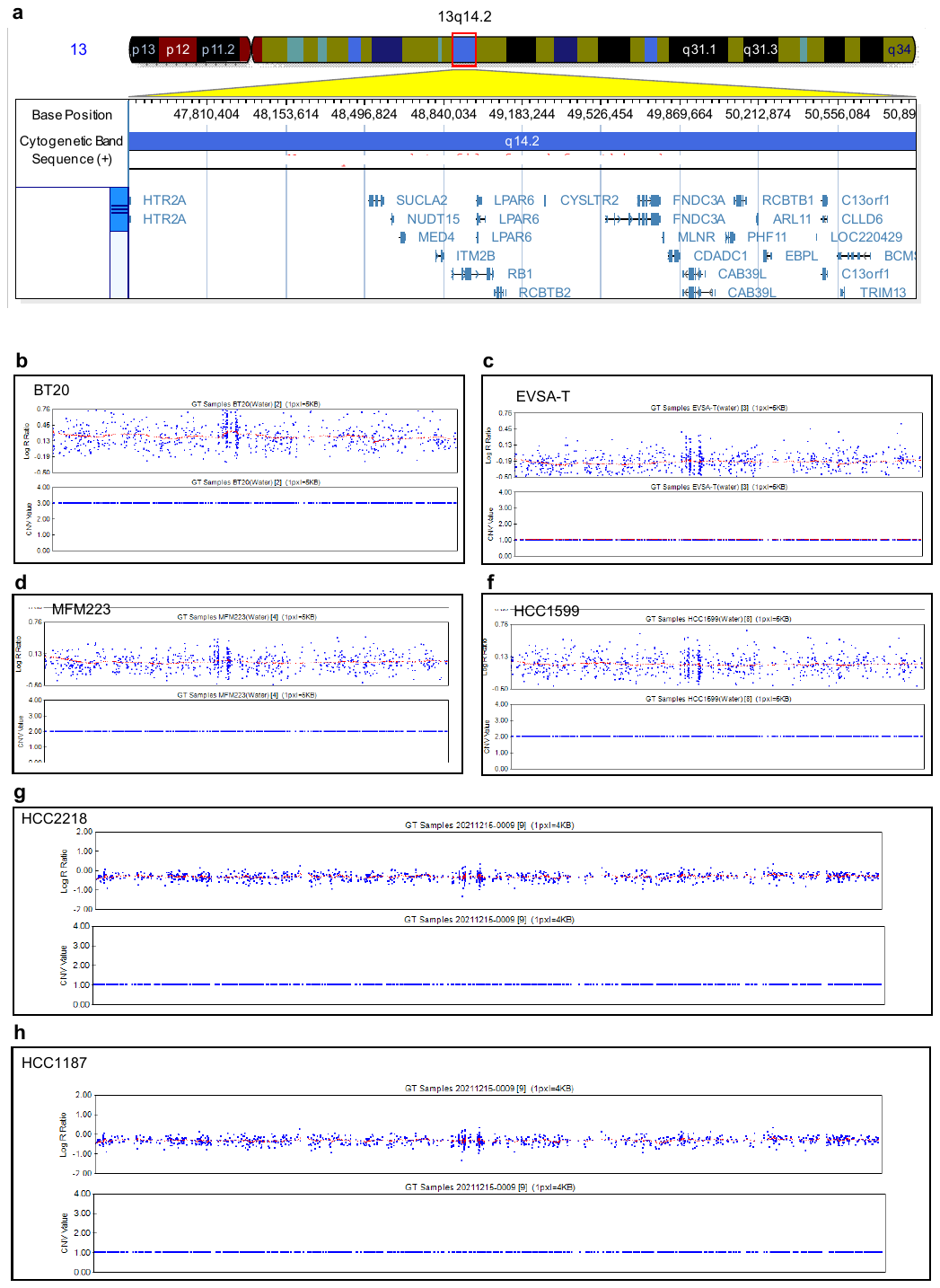

**Fig. S4.** Copy number profile by SNP6 analysis of breast cancer cell lines. **(a)** Schematic representation of chromosome 13q14.2 with the length in nucleotide as well as genes located in this region. **(b-h)** SNP6 analysis of DNA extracted from the breast cancer cell lines using genome studio. The plots indicate the copy number alteration of 13q14.2 cytoband. CNV Values: 1: loss, 2:diploid, 3: gain. data was processed with genome build ’hg19’.

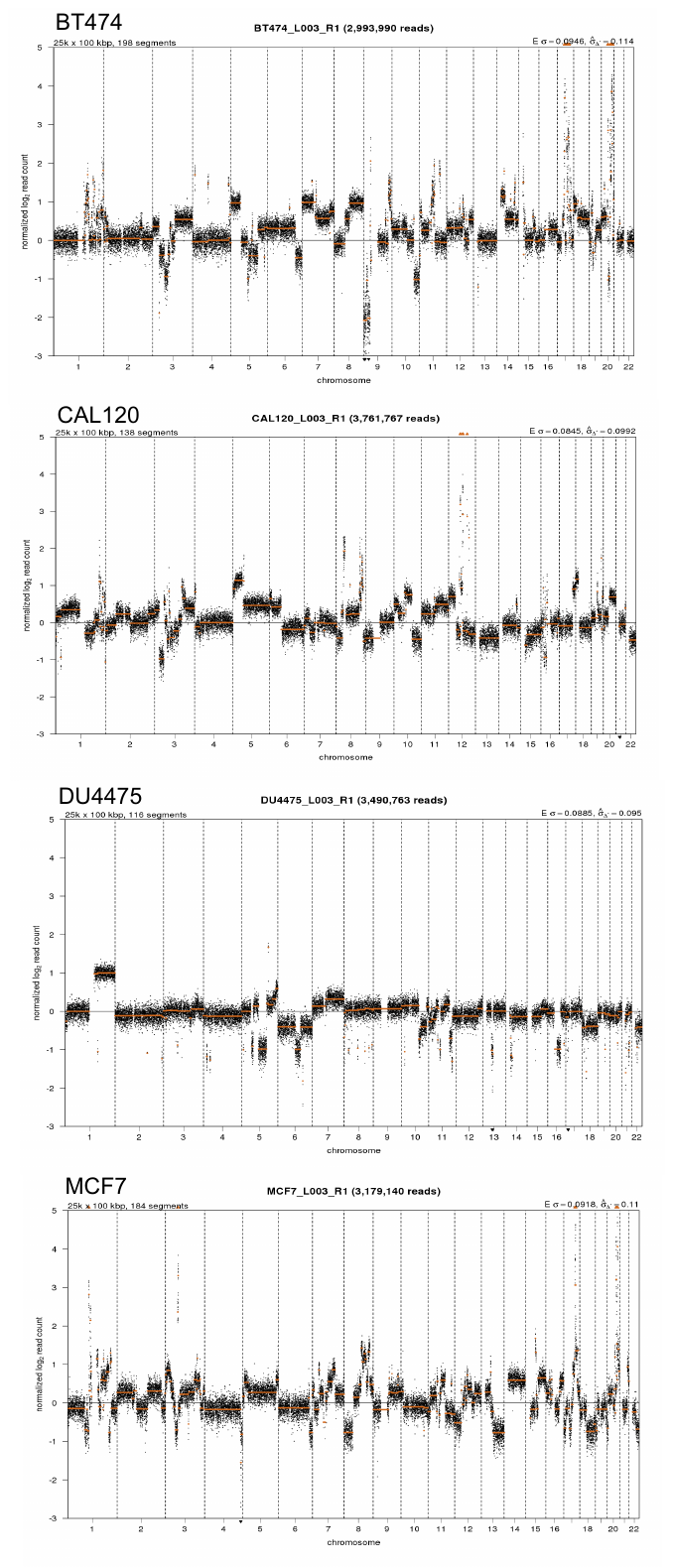

**Fig. S5.** Copy number profile by whole genome shallow sequencing analysis of breast cancer cell lines. Dots above the log2 ratio of 0 indicate regions of gains and amplifications and dots below log2 ratio of 0 indicate regions of losses and deletions. Data was processed with genome build ’hg19’.

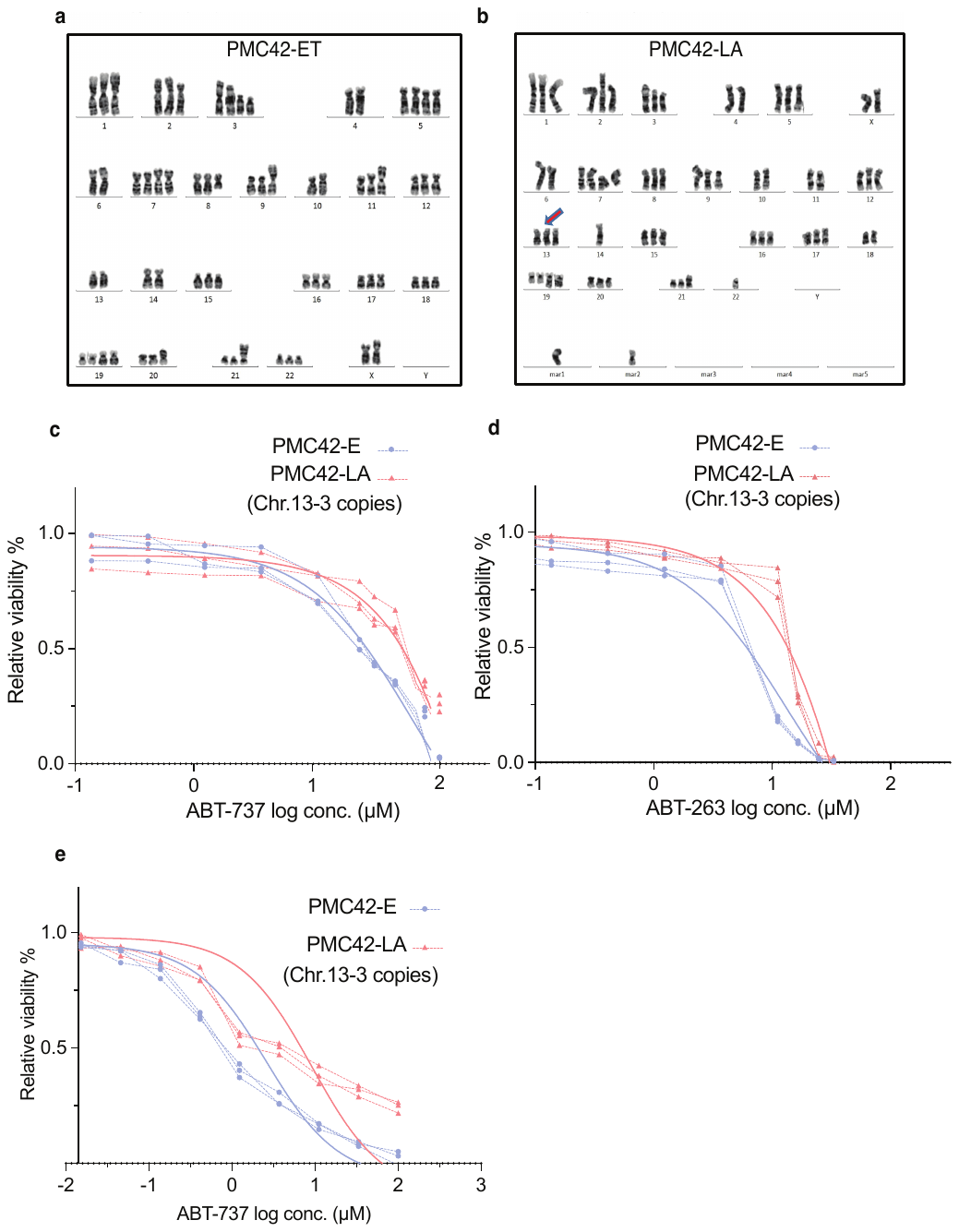

**Fig. S6.** *BCL2* drug response assessment to Chromosome 13q amplification. **(a)** G-banded karyotype of the triploid cell line PMC42-ET, reflecting aneuploidy with regards to numerical changes. **(b)** G-banded karyotype of the hyper diploid cell line PMC42-LA. Arrows point to the Chromosome 13 copy number difference. Mar chromosomes reflect derivative chromosomal with unknown origins. **(c-e)** IC50 measurement of log2 concentrations of ABT737, ABT263 and ABT199 in PMC42-ET and PMC42-LA cells after 72 hours treatment.

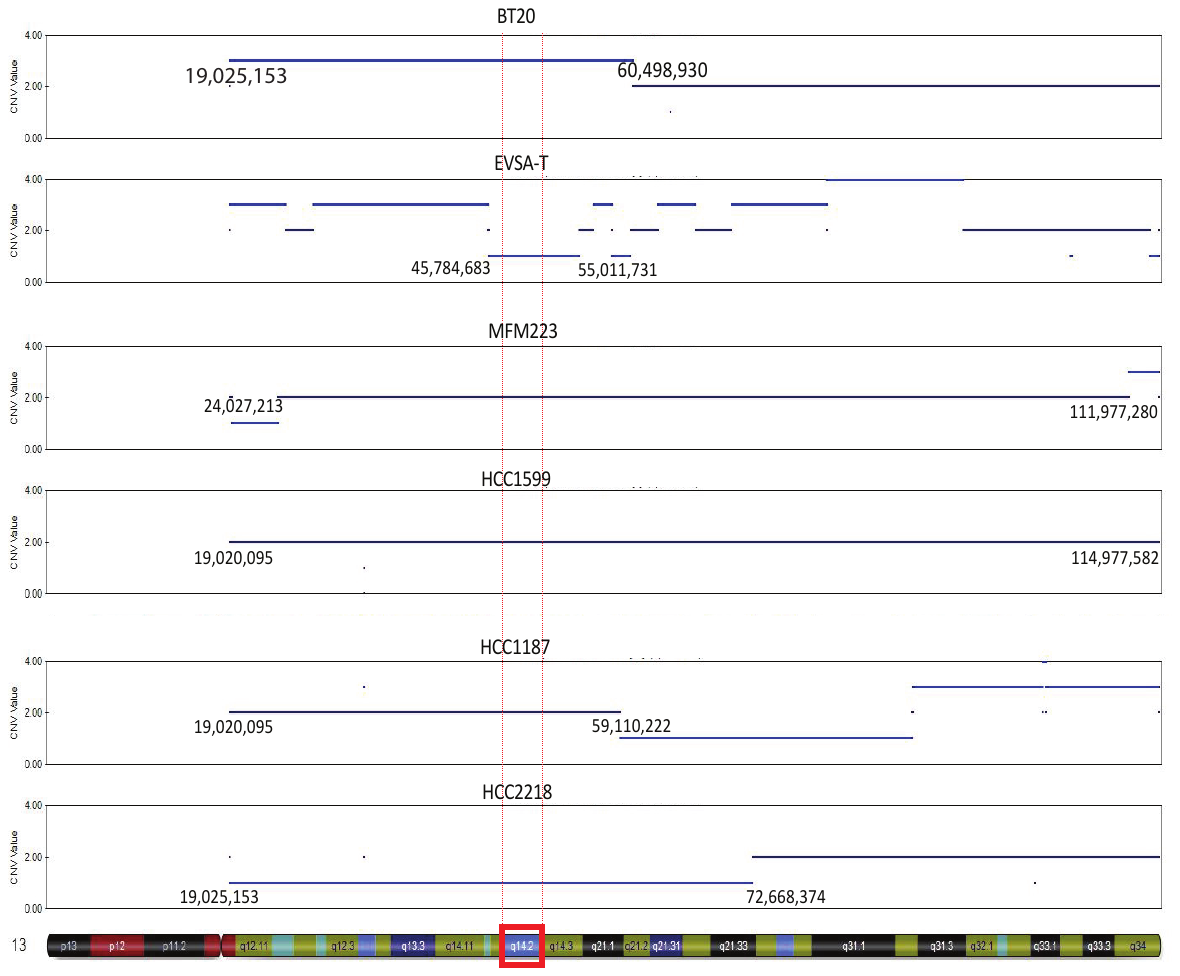

**Fig. S7.** Schematic representation of the length of the copy number alterations of chromosome 13q in breast cancer cell lines.
